# The triple-flash illusion reveals a driving role of alpha-band reverberations in visual perception

**DOI:** 10.1101/093500

**Authors:** Rasa Gulbinaite, Barkin İlhan, Rufin VanRullen

## Abstract

The modulatory role of spontaneous brain oscillations on perception of threshold-level stimuli is well established. Here, we provide evidence that alpha-band (7-14 Hz) oscillations not only modulate but also can drive perception. We used the “triple-flash” illusion: Occasional perception of three flashes when only two spatially-coincident veridical ones are presented, separated by ~100 ms. The illusion was proposed to result from superposition of two hypothetical oscillatory impulse response functions (IRF) generated in response to each flash (Bowen, 1989). In Experiment 1, we varied stimulus onset asynchrony (SOA) and validated Bowen's theory: the optimal SOA for illusion to occur was correlated, across subjects, with the subject-specific IRF period. Experiment 2 revealed that pre-stimulus parietal alpha EEG phase and power, as well as post-stimulus alpha phase-locking, together determine the occurrence of the illusion on a trial-by-trial basis. Thus, oscillatory reverberations create something out of nothing – a third flash where there are only two.

## INTRODUCTION

Spontaneous rhythmic fluctuations in various frequency bands have been consistently reported to affect perception across several sensory modalities (VanRullen, 2016b). Most studies found modulatory effects of pre-stimulus alpha-band (7-14 Hz) power, phase, and frequency on threshold-level stimulus detection, perceptual and temporal discrimination (Busch et al., 2009; Roberts et al., 2014; Baumgarten et al., 2015; Samaha and Postle, 2015). These perceptual consequences of endogenous oscillations imply that perception is inherently rhythmic and operates in a form of perceptual cycles, with periods of high and low excitability (Dugue et al., 2011; Vanrullen et al., 2011). However, most evidence for rhythmicity in perception is based on modulatory effects of ongoing oscillations on stimulus processing: For example, oscillatory phase at stimulus onset modulates stimulus visibility by approximately 10-15% (Mathewson et al., 2009; Busch and VanRullen, 2010; Dugue et al., 2011). Do brain oscillations merely modulate perception, or can they also, under certain conditions, drive perception over perceptual threshold and generate a percept without a physical stimulus?

In the “triple-flash” visual illusion, two brief light pulses separated by about 100 ms are sometimes perceived as three (Bowen, 1989). The effect was theoretically explained by the superposition of two damped oscillatory impulse response functions (IRF) generated in response to each stimulus flash: The third illusory flash is perceived when the delay between veridical flashes matches the period of oscillatory IRF. In this case, the later part of the oscillation is enhanced, and when this enhancement crosses perceptual threshold a third illusory flash is perceived (Fig. 1B, middle panel). However, when the delay does not match the period, the later part of the oscillation is dampened, and only two flashes are perceived (Fig. 1B, top and bottom panels). Thus, Bowen’s theoretical account of the triple-flash illusion assumes that the brain response to a single flash of light is oscillatory.

At first glance the triple-flash illusion appears similar to other phantom flash illusions (Wilson and Singer, 1981; Shams et al., 2000; Violentyev et al., 2005; Chatterjee et al., 2011; Apthorp et al., 2013), such as the sound-induced double-flash illusion, whose temporal window has been shown to be causally related to alpha-band oscillations (Cecere et al., 2015). However, there is one critical difference: The illusory third flash in the “triple-flash” illusion is purely endogenous, whereas perception of other phantom flash illusions requires simultaneous presentation of additional stimuli, either in a different modality or in a different spatial location.

Empirical tests of Bowen’s insightful predictions, and the role of oscillations in the generation of the triple-flash illusion, have not yet been demonstrated. Several findings, however, indicate that alpha-band oscillations could be involved in the generation of illusory third-flash percepts. First, the optimal delay of ~100 ms (or 10 Hz) between the two veridical flashes is within the frequency range of the endogenous alpha-band rhythm. Additionally, the optimal inter-flash delay for the illusion to be perceived varies across individuals (Bowen, 1989), and so does the alpha peak frequency. Differences in individual alpha peak frequency (IAF) have been reported both across individuals and across brain areas within individual (Doppelmayr et al., 1998; Haegens et al., 2014). Second, inter-individual variations in the frequency of occipital alpha oscillations are causally linked to the temporal properties of visual perception, such that faster occipital alpha oscillations are associated with finer temporal resolution in perception (Cecere et al., 2015; Samaha and Postle, 2015). Third, there is empirical evidence that the response to a single flash is indeed oscillatory, and reverberates at ~10 Hz up to a second after stimulus offset (VanRullen and Macdonald, 2012).

Understanding the role of oscillations in the triple-flash illusion is critical to the idea of perceptual cycles, as the illusory third-flash percept is potentially caused by perceptual reverberations – carryover effects of events unfolding *across* perceptual cycles. According to this line of reasoning, cortical excitability and stimulus-evoked responses determined by the power and phase of ongoing alpha-band oscillations at the moment of the first flash could have effects that last for several alpha cycles (Remond and Lesevre, 1967; Jansen and Brandt, 1991; Busch et al., 2009; Mathewson et al., 2009; Fiebelkorn et al., 2013). In Experiment 1, we directly tested whether illusory third-flash perception could result from summed reverberations of visual responses as proposed by Bowen (1989). In Experiment 2, we investigated prestimulus and stimulus-related effects of alpha phase, power, and frequency (separately for occipital and parietal alpha sources) on the perception of the illusory third flash.

## METHODS

### Experiment 1

This experiment consisted of two parts. In the first part, we estimated subject-specific optimal SOA of the triple-flash illusion using a psychophysical approach. In the second part, we established the subject-specific IRF using EEG recordings obtained during white noise flicker stimulation.

#### Participants

Thirty participants (12 females, mean age 27.4) took part in the psychophysics experiment. All participants had normal or corrected-to-normal vision. The data from three participants was excluded from the analyses: Two due to low accuracy on easy-to-discriminate long SOA three-flash trials, and one due to bias towards reporting illusions on very short SOA trials, with no discernable preferred SOA for the triple-flash illusion to be perceived. The study was conducted in accordance with the declaration of Helsinki and approved by the local ethics committee. Written informed consent was obtained from all participants prior to the experiment.

#### Stimuli and Procedure

Stimuli for the triple-flash experiment were two or three uniform white circles (radius 3.5°) presented on a black background in rapid succession peripherally above the fixation dot (eccentricity 7° visual angle). The viewing distance was constrained by the chin rest positioned 57 cm away from a 17-in. CRT monitor (600 × 480 pixels; 160 Hz vertical refresh rate). On each trial, either two or three circles were presented for a duration of one screen refresh (6.25 ms) with variable stimulus onset asynchrony (SOA; Fig. 1A). After the offset of the last flash, the screen remained black until a response was made. Following the response, the next trial started after a variable delay (1000–1500 ms). For a majority of participants (N = 20), the SOA on two-flash trials was randomly selected from one of 10 possible SOAs (50, 62.5, 75, 87.5, 100, 112.5, 125, 137.5, 150, 175 ms), and on three-flash trials from 8 possible SOAs (25, 31.25, 37.5, 50, 75, 87.5, 175, and 250 ms). For the remaining 10 participants, a finer sampling of SOAs was used: For the two-flash trials – 17 SOAs (50 to 250 ms, in steps of 12.5 ms), and for the three-flash trials – 27 SOAs (25 to 125 ms, in steps of 6.25 ms, and 125 to 250 ms, in steps of 12.5 ms). The overall probability of three flash trials was 33% throughout the experiment. The task consisted of 20 practice trials and 1200 experimental trials separated in blocks of 100 trials. Participants were tested in a dark room. They were instructed to keep their eyes on the fixation dot throughout the experiment, and to make speeded responses without sacrificing accuracy. “Left arrow” and “right arrow” keys were used to indicate perception of two- and three-flashes respectively. Participants were encouraged to take breaks after each block. The experiment lasted approximately an hour.

**Figure 1.**
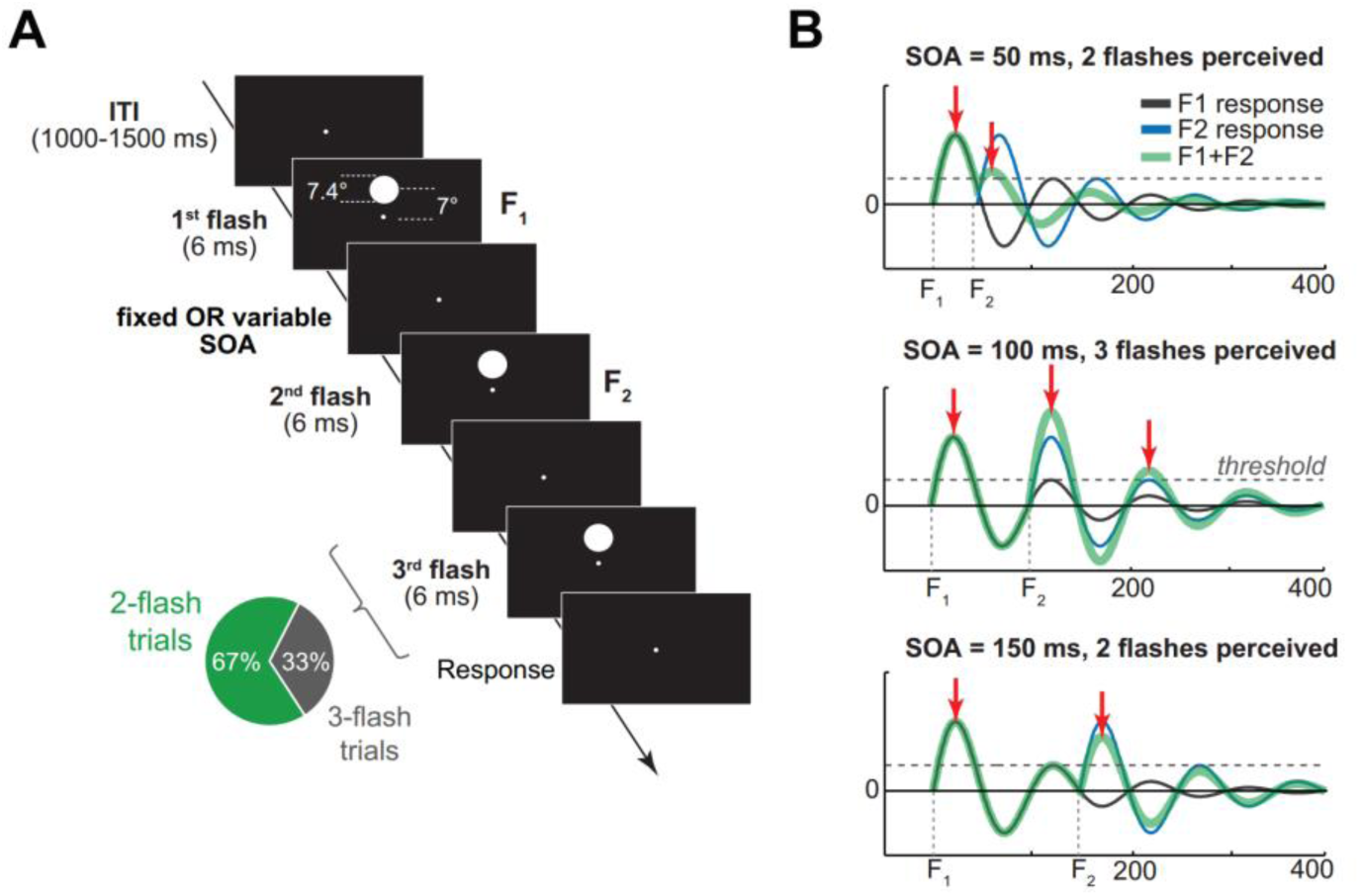
Trial structure and Bowen’s theoretical model. (A) On each trial two or three high-contrast circle stimuli were presented in rapid succession above the fixation dot. Stimulus onset asynchrony (SOA) on two-flash trials was either variable (Experiment 1), or fixed (Experiment 2); three-flash trial SOA was variable in both experiments.

In a separate experimental session, we recorded EEG while participants monitored a peripheral disk stimulus, whose luminance randomly varied every screen refresh with the constraint that the power spectrum of the luminance time course was flat between 0 and 80 Hz (i.e., white noise). Stimulus size, position, viewing distance, background, and screen refresh rate were kept identical to the triple-flash experimental session. Participants were instructed to detect target stimuli (a small circle surrounded by a darker annulus) presented in the center of the flickering stimulus by pressing a button. There were 2-4 targets presented during each 6.25 sec long trial. Target stimulus detectability (set to 50% using adaptive staircase procedure based on the performance during the first 30 trials) was manipulated by changing the relative contrast between the small circle and the annulus. The beginning of each trial was self-paced using a button press. The experiment lasted about one hour, and was divided in 8 blocks, with 48 trials in each block. Participants were encouraged to take rest breaks after each block.

#### Data analysis

To evaluate whether the triple-flash perception is related to oscillatory IRF, we correlated the period of subject-specific IRF with the subject-specific optimal SOA for illusory perception. Robustness of correlations was assessed by performing a bootstrapping procedure (resampling with replacement) using the Robust Correlation Toolbox (Pernet et al., 2012).

Subject-specific IRFs were obtained by cross-correlating time-series of stimulus luminance changes with concurrently recorded EEG time series at lags between −200 and 1500 ms (VanRullen and Macdonald, 2012). For this, EEG data were pre-processed using the same pipeline as for the data in the Experiment 2 (see below), with a few exceptions: (1) data were downsampled to 160 Hz to match the rate of stimulus luminance changes, (2) epochs were −250 – 6500 ms relative to the onset of the luminance sequence, (3) trials containing eye blink artefacts during luminance sequences were excluded from the analyses. The period of the individual IRF was estimated by performing the FFT of cross-correlation result, finding the peak frequency (*f*) in the range of 6-14 Hz, and expressing it as period in milliseconds (1/*f*).

Subject-specific optimal SOA for the triple-flash illusion to be perceived was determined by fitting symmetrical and non-symmetrical functions to the behavioral performance on two-flash trials (proportion of two-flash trials perceived as three). Initial to-be-fitted model parameters were based on each subject's empirical data, and used as an input for *fminsearch* Matlab function. We used Gaussian (1), Weibull (2), and ex-Gaussian (3) functions:

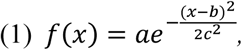

where *a* is Gaussian peak y-axis value (initially set to maximum number of illusions perceived by participant); *b* Gausian peak x-axis value (initially set to SOA of maximum number of illusions); *c* is width of the Gaussian (initially set to 0.05 sec).

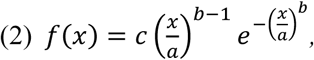

where *a* is the x-axis scaling factor (initially set to SOA of maximum number of illusions perceived by participant); *b* defines the shape of the curve (initially set to 4); and *c* scales the curve along the y-axis (initially set to twice the maximum number of illusions).

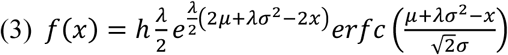

where λ = 1/τ, and τ is an exponential decay parameter (initially set to 0.1); *μ* is mean of the Gaussian (initially set to half the SOA with maximum number of illusions); *σ* is variance of the Gaussian (initially set to 0.05 sec); *h* is the y-axis scaling factor (initially set to maximum number of illusions divided by four).

Goodness of fit at each *fminsearch* iteration and across the three different fitting functions was evaluated using R^2^, which is the amount of variance accounted for.

### EXPERIMENT 2

The purpose of Experiment 2 was to characterize EEG changes that accompany perception of the third-flash illusion relative to no-illusion trials with physically identical stimuli.

#### Participants

EEG data were recorded from 35 participants (18 females, mean age 26.5), 27 of them also participated in Experiment 1. All participants had normal or corrected-to-normal vision. The study was conducted in accordance with the declaration of Helsinki and approved by the local ethics committee. Written informed consent was obtained from all participants prior to the experiment.

#### Stimuli and Procedure

In comparison to the procedure of Experiment 1, two-flash trial SOA was fixed at 87.5 ms. The choice of this SOA was based on the results of Experiment 1, where maximal number of illusions on average across participants based on ex-Gaussian fits was 92 ms (*SD* =17 ms). As in Experiment 1, three-flash trial SOA was variable (25, 31.25, 37.5, 50, 75, 87.5, 175, and 250 ms), and overall proportion of three-flash trials was kept at 33% throughout the experiment. To avoid muscle artifacts in EEG, viewing distance was unconstrained by the chin rest but still kept at approximately 57 cm.

#### Data acquisition and preprocessing

EEG data were recorded using 64-channel ActiveTwo BioSemi system (for detailed description of the hardware see www.biosemi.com) at 1024 Hz sampling rate. Offline, the data were down-sampled to 512 Hz and re-referenced to the average activity over all electrodes. Continuous EEG recordings were band-pass filtered at 0.5 – 80 Hz, and electrical line noise was removed using a notch filter (band-stop 47 to 53 Hz). The data were then epoched (-1500 ms to 2000 ms relative to the first stimulus onset), and baseline-corrected with respect to the time window of −200-0 ms relative to the first stimulus onset.

Artifact removal was done in two steps. First, the data was visually inspected and trials containing muscle artifacts or eye blinks during and 800 ms prior to the stimulus presentation were removed. The second artifact rejection step involved independent components analysis (ICA; Delorme and Makeig, 2004). Components that did not account for any brain activity, such as eye-movements or noise, were subtracted from the data (on average, 1.4 components per subject). Furthermore, trials with reaction time (RT) of 3 standard deviations longer than subject’s mean RT were excluded from the analysis. On average, 90.6 % of two-flash trials per subject were included in the analysis (SD = 4.4%).

#### Alpha-band source separation

To account for variability in alpha peak frequency across individuals, and across brain regions within an individual (Haegens et al., 2014), we separated parietal and occipital alpha sources based on ICA using JADE algorithm as implemented in EEGLAB and dipole fitting using DIPFIT toolbox (Oostenvelt et al., 2003; Delorme and Makeig, 2004). First, eye-blink and movement artifact-free data was band-pass filtered at 5-15 Hz. Thereafter, ICA was performed on pre-stimulus time window (-1000 to 0 ms, where 0 is the first stimulus onset). For each independent component (IC) map that had a clear alpha peak in the frequency spectrum, a current dipole model was fitted using a three-shell boundary element model. One parietal and one occipital IC was selected based on the spatial proximity of the fitted dipoles to the reference anatomical coordinates, with constraints that selected equivalent dipoles had less than 15% residual variance from the spherical forward-model scalp projection, and were located inside the model brain volume. Reference anatomical coordinates for parietal ROIs were centered on the left and on the right Brodmann area 7 (right-side MNI coordinates: −20 −90 0; left-side MNI coordinates: 20 −90 0); for occipital ROIs, mean coordinates of Brodmann areas 17 and 18 were used (right-side MNI coordinates: −20 −70 50; left-side MNI coordinates: 20 −70 50; Haegens et al., 2014). Finally, individual alpha-peak frequency (IAF) at occipital and parietal sources was estimated by taking the FFT of IC time series, and finding the peak in the range of 6–14 Hz.

#### Independent component time-frequency analyses

Epoched unfiltered data was multiplied by ICA unmixing matrix from selected subject-specific parietal and occipital components to obtain component single-trial time series. Time-frequency decomposition was performed by convolving single-trial data from parietal and occipital ICs with complex Morlet wavelets, defined as:

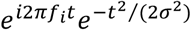

where *t* is time, *f_i_* is frequency which ranged from 2 to 40 Hz in 39 logarithmically spaced steps, and *σ* is the width of each frequency band defined as *n*/(2*πfi*), where *n* is a number of wavelet cycles that varied from 1 to 7 in logarithmically spaced steps to obtain comparable frequency precision at low and high frequencies.

Instantaneous power was computed as the square of the analytic signal Z (a complex result of convolution) and averaged across trials (i.e., *power =* Re[Z(t)]^2^ + Im[Z(t)]^2^). Thus obtained power values were then baseline-corrected by converting to decibel scale (10 log_10_(*power/baseline*)), where condition-average power from −400 to −100 ms pre-stimulus period served as the baseline. Condition-average rather than condition-specific baseline was used to avoid introducing spurious power differences in the post-stimulus period.

To evaluate the effects of alpha phase on illusory third-flash perception, we compared phase distributions on illusion vs. non-illusion two-flash trials using phase opposition sum (POS), which is expressed as a function of inter-trial phase clustering (ITPC) of illusion and non-illusion trials relative to ITPC of all trials (VanRullen, 2016a):

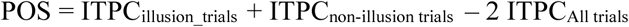

where 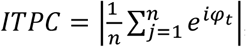, the phase angle at each time point φ_t_= arctan (Im[Z(t)]/Re[Z(t)]), *n* is the number of trials, *j* is the trial, and *i* is the complex operator.

To test the hypothesis that third-flash perception is related to more precise phase alignment of oscillatory responses to veridical flashes, we computed phase consistency across trials at 11.43 Hz (corresponding to the 87.5 ms SOA) using the weighted pair-wise phase consistency (wPPC) metric (Vinck et al., 2010). Consistency of phases across trials is often estimated using ITPC. However, ITPC is sensitive to relative and absolute trial count (Cohen, 2014a), whereas wPPC corrects for this bias because it measures similarity of phase angles between all trial pairs. The formula for wPPC as implemented in Fieldtrip function *ft_connectivity_ppc.m* is:

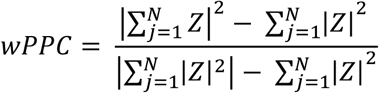

where Z is a complex convolution result computed at each time point using 3-cycle Morlet wavelet as described above. Conceptually, wPPC measures the extent to which the circular distance between phase angle pairs taken from different trials is non-uniformly distributed, whereas ITPC measures the extent to which a distribution of phase angles across trials at each time point is non-uniform. Analytically, wPPC is comparable to ITPC squared (Vinck et al., 2010).

#### Statistical analyses

Differences in the pre-stimulus and post-stimulus alpha-band power were performed taking into account individual alpha peak frequency (IAF), and its variability across parietal and occipital alpha sources (Haegens et al., 2014). Therefore, for each participant alpha band was defined as IAF±1.5 Hz separately for parietal and occipital alpha sources. Statistical comparison of pre-stimulus alpha-band power was performed on raw, i.e. non-baseline-corrected power values. Statistical comparison of pre-stimulus (-600 − 0 ms) and post-stimulus (0 − 600 ms) alpha-band time courses, and wPPC between illusion and non-illusion trials was performed using non-parametric permutation testing procedure as described further (Maris and Oostenveld, 2007).

First, one-sample t-tests (separate for parietal and occipital alpha components) were performed comparing time-series of condition difference in power (or wPPC) against 0 (real t-value). Second, a null hypothesis distribution was created by multiplying the condition difference time-series from a random number of participants by -1. This procedure was repeated 1000 times to obtain surrogate t-values. Third, power (or wPPC) differences were considered statistically significant if real t-value at that time point was larger than 95% of surrogate t-values. Finally, cluster-based correction was applied to correct for multiple comparisons over time points. Clusters of contiguous time points were considered significant if the size of the cluster (sum of t-values within a cluster) was bigger than expected under the null hypothesis at a significance level of p < .05. The null hypothesis distribution of cluster sizes was obtained by thresholding the surrogate t-values at p < .05, and storing the maximum cluster size values. Finally, to obtain more stable estimates from permutation testing we ran a “meta-permutation test” by repeating the permutation procedure 20 times. Thus, averaged results from 20 meta-permutations, each consisting of 1000 iterations, are reported.

Statistical significance of POS was tested by comparing the observed POS value at each time-frequency point to the null hypothesis distribution of POS values, which was obtained using the following procedure. For each participant, illusion and non-illusion trials were randomly relabeled and surrogate POS was computed. This was repeated 1000 times at each time-frequency tile. Thus obtained surrogate POS values were then used to compute 100.000 grand-average surrogate POS values by randomly selecting one of the surrogate POS values for each participant. Finally, p-value was computed as the proportion of grand-average surrogate POS values that exceeded empirically observed grand-average POS. This p-value indicates how empirically observed phase differences between illusion and non-illusion trials deviated from phase differences expected under null hypothesis (Fig. 6). Statistical comparison of POS frequency profile was performed by summing the observed and surrogate grand-average POS values over time (−600 to 0 ms) and computing the p-value as described above. Correction for multiple comparison across frequencies was performed using non-parametric permutation testing procedure, equivalent to that used for alpha power and wPPC comparison between conditions.

## RESULTS

### Experiment 1

In Experiment 1, we empirically tested Bowen’s theoretical idea that the triple-flash illusion occurs when the delay between veridical flashes matches the period of IRF generated in response to a single flash. The two key components of Bowen’s model were determined using a psychophysical approach (subject-specific optimal delay between flashes) and EEG recordings (period of the subject-specific IRF).

### Evidence for Bowen’s model

Replicating Bowen’s results (1989), we found that illusory third flash perception depended on SOA between the two veridical flashes. The main effect of SOA on two-flash trials was significant for both coarsely (*F*(1,9) = 20.86, p < .001) and finely sampled SOAs (*F*(1,16) = 4.85, p < .001), such that perception of illusions followed an inverted u-shape function (Fig. 3A). Averaged across finely and coarsely sampled equivalent SOAs, a 75 ms delay between two veridical flashes yielded the strongest third-flash illusion: At this SOA, the illusory third flash was perceived on about half of the two-flash trials (*M* = 41.41%, *SD* = 26.83%). In contrast, three veridical flashes separated by the same 75 ms SOA almost always were perceived as three (*M* = 91.04%; *SD =* 11.86%), indicating that perception of the illusory flash on two-flash trials does not result from an inability to distinguish rapidly presented stimuli. This was further supported by the main effect of SOA on three-flash trials (for finely sampled SOAs *F*(1,26) = 23.27, p < .001; for coarsely sampled SOAs *F*(1,7) = 94.35, p < .001), where perception of three flashes steadily increased as a function of SOA (Fig. 3A).

**Figure 3.**
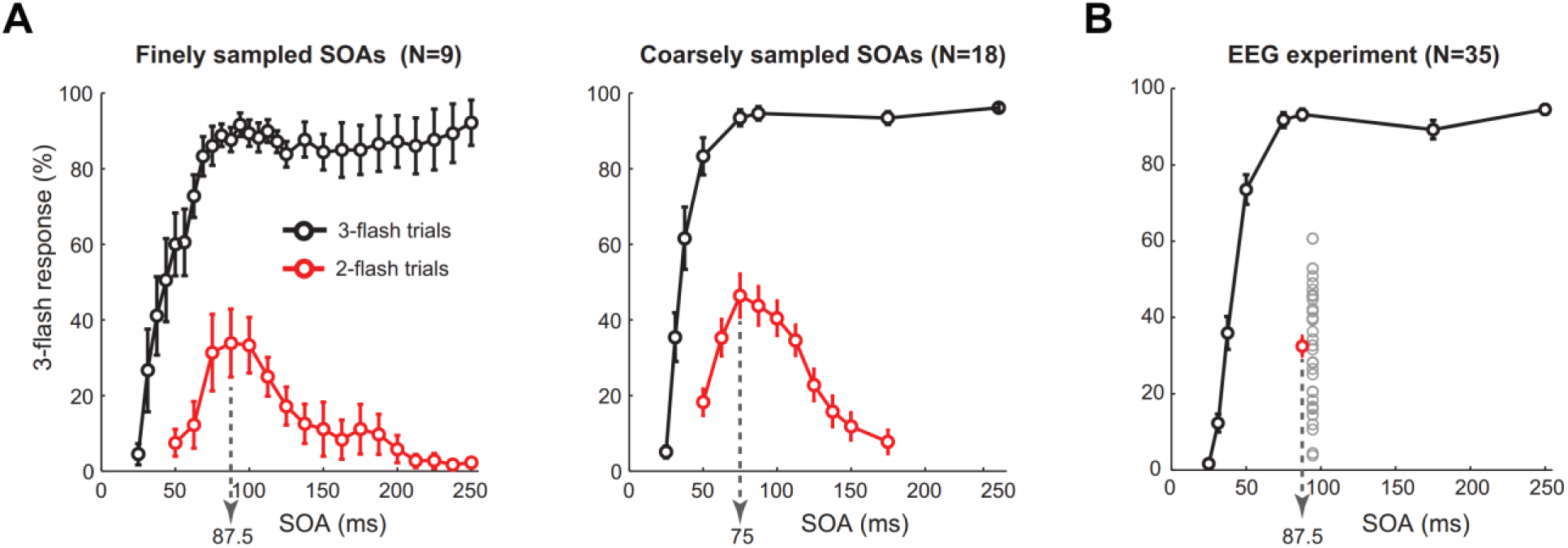
Behavioral performance. Percentage of three-flash percepts for two- and three-flash trials as a function of stimulus onset asynchrony (SOA) between veridical flashes. (A) Behavioral results of psychophysics Experiment 1, where fine (17 SOAs for two- and 27 SOAs for three-flash trials) and coarse (10 SOAs for the two- and 8 SOAs for the three-flash trials) were used. Dotted arrows indicate two-flash trial SOA that produced on average the maximum number of illusions. (B) Behavioral results of the EEG experiment (Experiment 2), where the two-flash trial SOA was fixed at 87.5 ms. Grey circles represent average percent of illusions for each participant (N=35). Error bars show standard error of the mean.

To directly test Bowen’s theoretical account of the triple-flash illusion, we first determined each participant’s impulse response function (IRF) and its period. As previously reported (VanRullen and Macdonald, 2012), white noise stimuli that have a flat frequency spectrum can be used to reveal subject-specific IRF by cross-correlating stimulus luminance time series with concurrently recorded EEG time series (Fig. 4B). As in previous reports (Ilhan and VanRullen, 2012; VanRullen and Macdonald, 2012), the cross-correlation result was oscillatory, with a period of ~100 ms on average and maximal amplitude over occipital electrodes (Fig. 4D). Next, we determined subject-specific SOA that maximized illusory third-flash perception by fitting symmetrical and non-symmetrical functions (Gaussian, Weibull, and exGaussian) to behavioral performance on two-flash trials. Given that exGaussian function provided the best model fits *(M(R^2^_Gaussian_)* = 0.89, *SD(R^2^_Gaussian_)* = 0.09; *M(R^2^_Weibull_)* = 0.89, *SD(R^2^_Weibull_)* = 0.08; *M(R^2^_exGaussian_)* = 0.91, *SD(R^2^_exGaussian_)* = 0.08), the peak of fitted exGaussian function was taken as a subject-specific optimal SOA (Fig. 4A).

The correlation between two key elements of Bowen’s model – subject-specific SOA that maximized illusory perception and period of subject-specific IRF – was significant (*r*_Pearson_ (25) = 0.51, *p* = .003, CI = [0.17 0.75]; Fig. 4C). This result is the first direct evidence for Bowen’s theoretical account of the triple-flash illusion. To evaluate the influence of the outliers, we recomputed correlation for 1000 times by randomly resampling with replacement. The 95% percentile CI of these bootstrapped correlations did not include 0 (CI_95%_ = [0.17 0.72]), indicating robustness of the observed effect. For consistency purposes with other analyses, we also tested resistance to outliers by using a non-parametric permutation testing procedure, by randomly shuffling IRF values across participants and recomputing correlation for 1000 times (Maris and Oostenveld, 2007). The empirically observed correlation was significantly different from the null hypothesis distribution (z-score = 2.69, p = .0036), indicating robustness of the observed effect.

### Experiment 2

In Experiment 2, we investigated the role of alpha-band oscillations in generation of the triple-flash illusion. For this we conducted an EEG experiment, for which SOA on two-flash trials was fixed at 87.5 ms based on the results of Experiment 1, where maximal number of illusions on average across participants for ex-Gaussian fits was 92 ms (*SD* = 17 ms). The fixed SOA was chosen to maximize the number of illusion trials for subsequent phase-based analyses that are sensitive to the number of trials (Vinck et al., 2010; Cohen, 2014a; VanRullen, 2016a).

**Figure 4.**
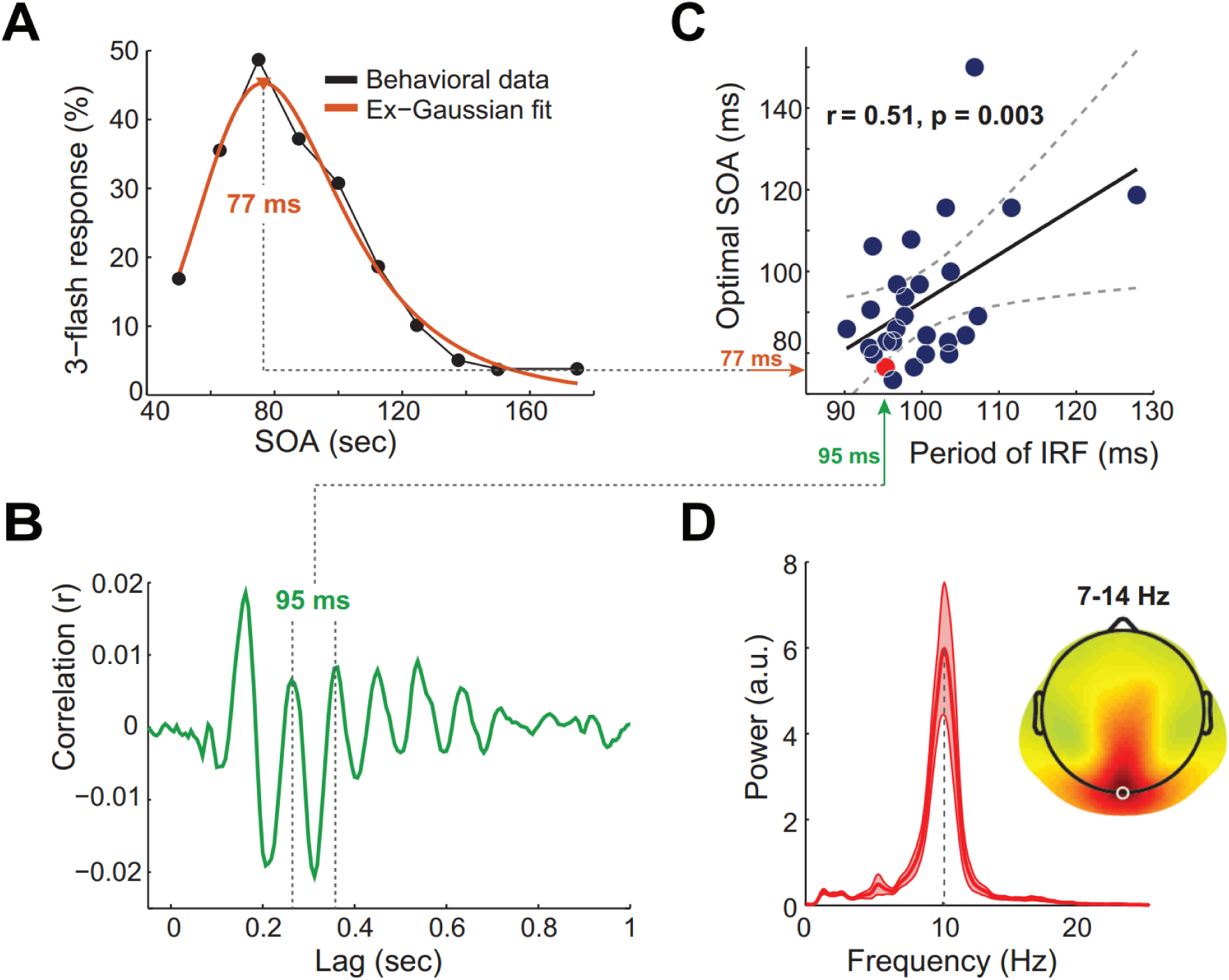
Relationship between period of impulse response function (IRF) and optimal delay between flashes. (A) Single-subject behavioral data showing the probability of third-flash perception on two-flash trials as a function of SOA (black line) and exGaussian fit used to determine subject-specific optimal SOA (orange line). (B) Single-subject IRF at Oz electrode (same participant as in panel A). (C) Correlation (across participants, N=27) between period of IRF at Oz electrode and optimal SOA. Dashed curves represent 95% confidence intervals around the slope of regression line. Red dot indicates participant’s data depicted in panels A and B. (D) Subject-average power spectrum of IRF at Oz and topographical map of 7-14 Hz power indicating that IRF is strongest around Oz (white circle). Subject-average power spectrum of IRF at Oz electrode with a peak centered at 10 Hz. Light red areas represent standard error of the mean.

### Third-flash perception depends on individual alpha frequency

Why did some participants perceive the illusion nearly half of the time and others perceived virtually none (Figure 3B)? Based on the previously reported correlation between IRF frequency and occipital individual alpha-peak frequency (IAF; VanRullen and Macdonald, 2012), we hypothesized that variability in IAF could be related to the observed between-subject differences in proneness to perceive the third-flash illusion. Specifically, the closer the match between IAF and fixed SOA used in the EEG experiment (87.5 ms ~ 11.43 Hz), the more illusions a given participant would perceive. We focused on the frequency of task-related (as opposed to resting-state) alpha-band oscillations prior to the first flash, because IAF has been shown to be state dependent (Haegens et al., 2014). Furthermore, parietal and occipital IAFs were determined in the pre-stimulus window to avoid contamination from sensory stimulus processing that is accompanied by rapid instantaneous frequency changes in all frequency bands (Burgess, 2012). To account for IAF variability across brain regions (Klimesch, 1999; Haegens et al., 2014), we isolated occipital and parietal alpha sources using independent component analysis (for details, see Methods section). For two participants, alpha-band sources could not be determined due to small alpha peaks in the power spectrum that were indistinguishable from noise. Thus 33 participants were included in the correlation analyses. Average peak frequency in parietal ROI was 10.4 Hz (SD = 0.94), and in occipital ROI – 10.5 Hz (SD = 1.1). Average frequency of alpha oscillations across participants statistically did not differ between the two regions of interest (*t*(32) = −0.625, *p* = .536).

The probability of the third-flash perception using fixed SOA was correlated with IAF at parietal alpha sources (*r*_Pearson_ (31) = −0.52, *p* = .002, CI = [−0.73 −0.21]; Fig. 5C): The smaller the absolute distance between parietal IAF and 11.43 Hz, the more illusions participant perceived. The same correlation for IAF at occipital sources was not significant (*r*_Pearson_ (31) = −0.33, *p* = .063, CI = [-0.60 0.02]; Fig. 5D). Robustness of the correlations was evaluated using bootstrapping procedure, which revealed that the 95% percentile CI of bootstrapped correlations for parietal sources did not include 0 (CI = [−0.74 −0.20]), whereas for occipital sources it did (CI = [−0.62 0.14]). A non-parametric permutation testing procedure indicated that the empirically observed correlation was significantly different from the null hypothesis distribution (z-score = −2.97, p = .0015) for the parietal as well as occipital IAF (z-score = −1.85, p = .03).

It is important to note that IAF is not stationary and fluctuations around IAF are relevant for perception (Cohen, 2014b; Samaha and Postle, 2015). However, momentary changes in IAF are an order of magnitude smaller (~ 0.04 Hz, see Samaha and Postle, 2015) than IAF differences between parietal and occipital ROIs when compared across subjects (0.6 Hz).

**Figure 5.**
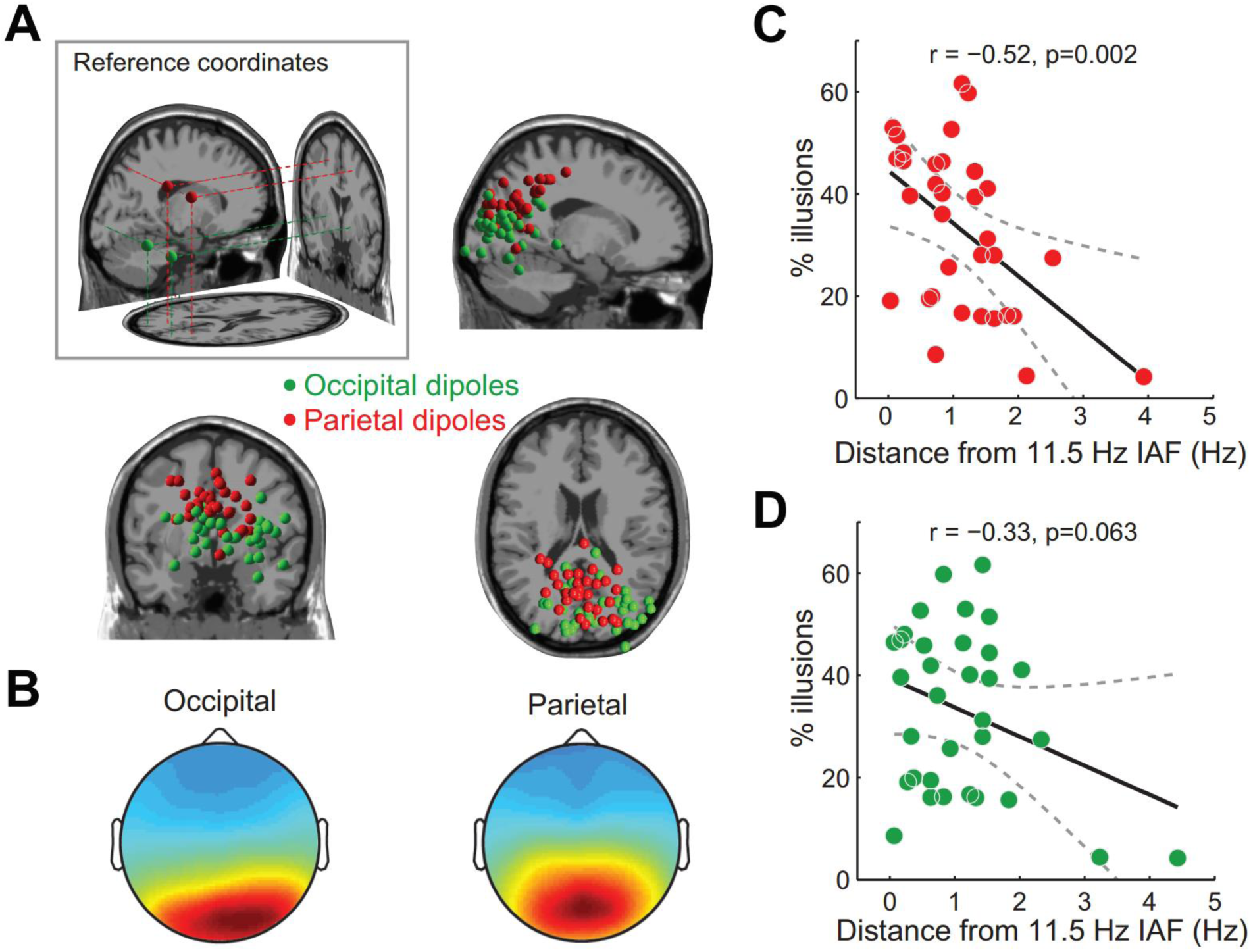
Relationship between the frequency of alpha oscillations and third-flash illusory percepts. (A) Dipole locations for anatomical reference coordinates for parietal and occipital ROIs (left-most panel). Locations of equivalent dipoles for parietal and occipital alpha independent components. (C, D) Correlations (across participants, N=33) between individual alpha peak frequency (IAF) and overall percent of illusions, indicating that the proportion of illusions depended on the match between inter-flash delay (11.43 Hz ~ 87.5 ms) and IAF at parietal (C) but not occipital alpha sources (D). (B) Scalp projection of cluster centroids for occipital and parietal independent component (IC) clusters, revealing more central and anterior projection for parietal ICs.

### Pre-stimulus alpha phase predicts third-flash perception

Although correlation between subject-specific optimal SOA and the period of subject-specific IRF lends support to Bowen’s notion that the triple-flash illusion reflects a superposition of two oscillatory responses, it does not explain trial-to-trial variability in perception of the illusion. At the subject-specific optimal SOA, the third flash is only perceived on average half of the time (45% in the Experiment 2). To address this question, we contrasted brain activity during physically identical two-flash trials on which the third-flash was either reported, or not.

We examined the effects of pre-stimulus alpha phase on perceptual outcome by computing the phase opposition sum (POS), a measure that represents the extent to which phase distributions between two classes of trials differ (here, illusion and non-illusion trials; Busch et al., 2009; VanRullen, 2016a). To minimize the effects of unequal illusion and non-illusion trial counts on the reliability of POS – an inherent feature of all phase-based time-frequency analyses methods (Vinck et al., 2010; Cohen, 2014a) – we selected only participants for which illusion and non-illusion trial counts differed by less than 10% (N=14). As for between-subject analyses, within-subject analyses were performed separately for occipital and parietal alpha sources.

If the phase of spontaneous alpha oscillations prior to the first flash is predictive of the third-flash perception, we should observe a strong phase clustering around a certain phase angle for illusion trials accompanied by strong phase clustering around the opposite phase angle for non-illusion trials. Prestimulus alpha phase differed at parietal but not occipital alpha sources (Fig. 6): The POI spectrum (averaged across all time points in the pre-stimulus interval) at parietal alpha sources was statistically significant in 6-12 Hz frequency range (corrected for multiple comparisons across frequencies using cluster-based permutation testing). Although analyses were performed on alpha sources, finding phase-opposition in the alpha band is not trivial, as source-separation was not based on phase measures.

**Figure 6.**
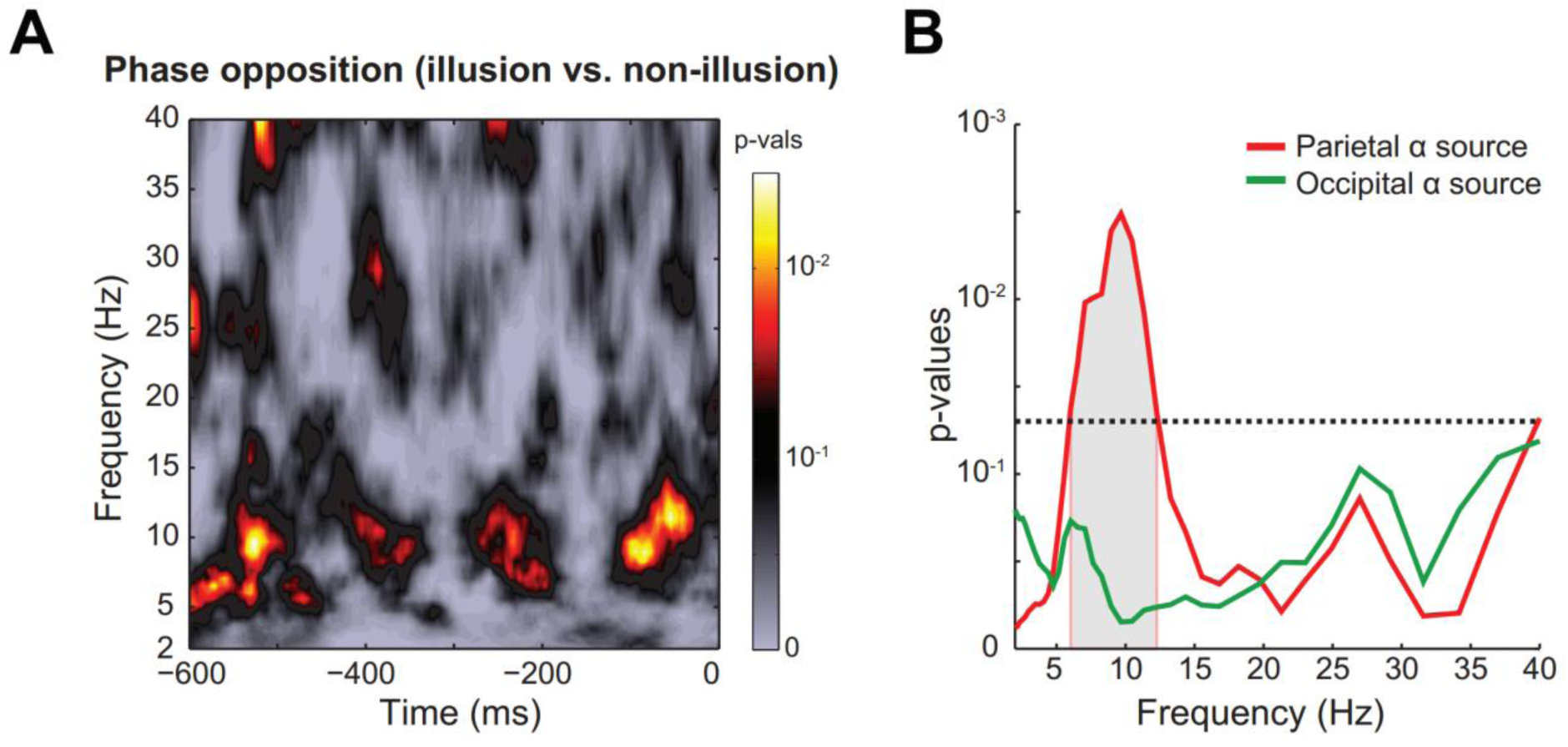
Pre-stimulus alpha phase differences between illusion and non-illusion trials. (A) Time-frequency representation of p-values (for parietal alpha sources) computed as a proportion of surrogate phase opposition values (distribution of phase opposition values expected under null hypothesis) that exceeded empirically observed phase opposition values. (B) Frequency profile of phase opposition (averaged over −600 – 0 ms pre-stimulus time window). Grey shaded area represents frequencies at which the observed phase opposition frequency profile was significantly different from the frequency profile of surrogate phase opposition values (corrected for multiple comparisons across frequencies using cluster-based permutation testing). The p-value of 0.05 is marked by a horizontal line.

Next we tested the effects of pre-stimulus alpha power, which has been shown to affect both near-threshold stimulus perception as well as perception of illusions (Romei et al., 2008; Lange et al., 2014). Within-subject analysis revealed significantly lower pre-stimulus alpha power on illusion vs. non-illusion trials at parietal but not occipital alpha sources (Fig. 7; all p values < 0.05 in the time interval -520 – -270 ms relative to first flash, corrected for multiple comparisons using cluster-based permutation testing).

**Figure 7.**
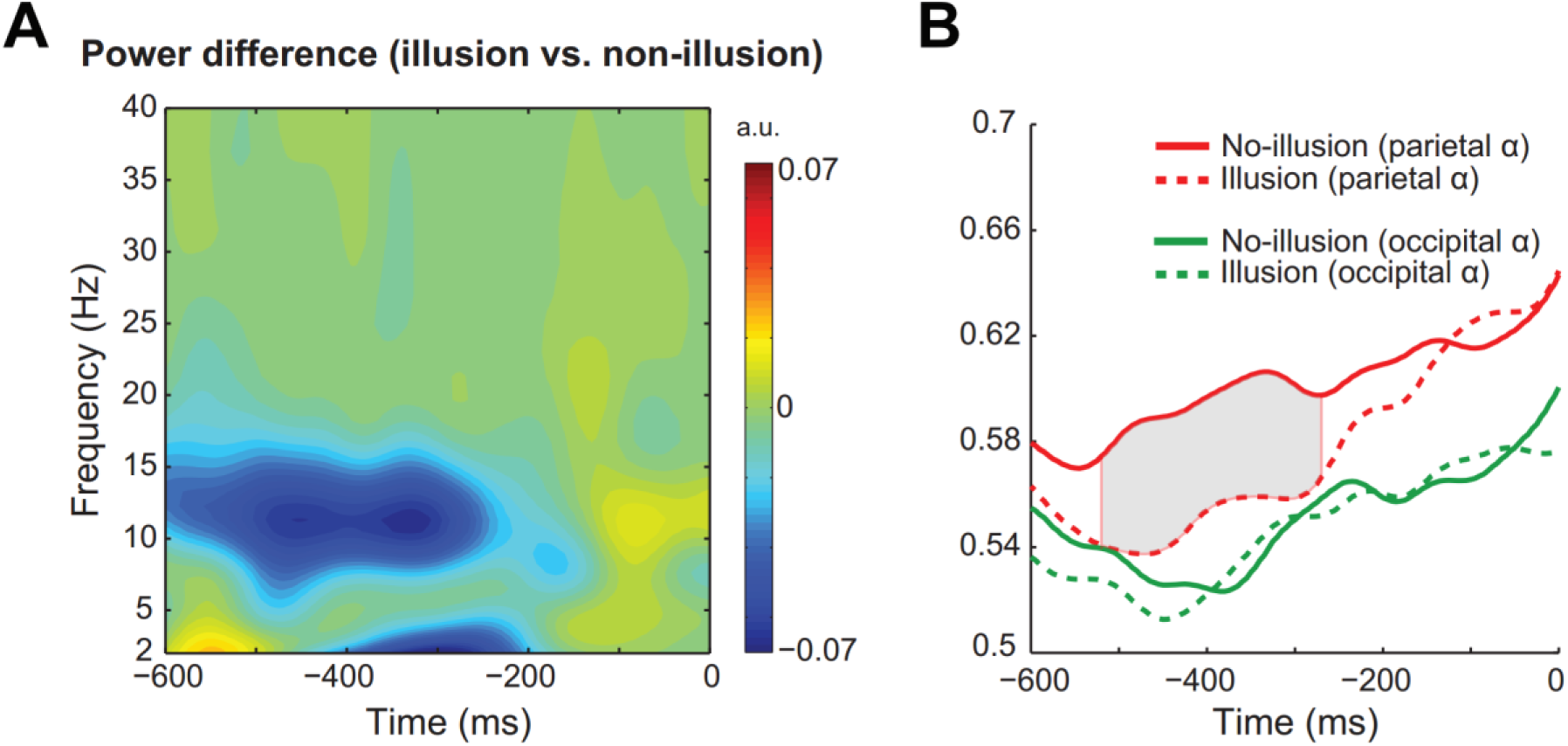
Pre-stimulus alpha power differences between illusion and non-illusion trials. (A) Time-frequency representation of alpha power differences at parietal sources. (B) Time-courses of alpha power (IAF±1.5 Hz) for illusion (dashed lines) and non-illusion (solid lines) trials at occipital and parietal alpha sources, demonstrating that alpha power differences were present at parietal but not occipital alpha sources. Grey shaded area represents the time interval where statistically significant differences between the two trial groups was observed (corrected for multiple comparisons using cluster-based permutation testing).

### Illusory perception is associated with stronger post-stimulus local phase realignment

According to Bowen’s theoretical model, presentation of the second flash in-phase with the oscillatory IRF evoked by the first flash would result in perfect superposition of the two IRFs and hence the perception of an illusory third flash. Whenever, for any reason (e.g. non-optimal SOA, non-optimal alpha phase at the first-flash onset, variability in stimulus-evoked oscillatory alpha phase, etc.), the second flash does not occur in-phase with the oscillatory response evoked by the first flash, then the model stipulates that third-flash perception would be less likely. Thus, we predicted more precise phase alignment on illusion than non-illusion trials in the post-stimulus window. We assessed phase alignment by computing weighted pair-wise phase consistency metric (wPPC; Vinck et al., 2010) at the frequency of the two veridical flashes (11.43 Hz ~ 87.5 ms). As illustrated in Figure 8, we found significantly higher wPPC for illusion than for non-illusion trials at parietal but not occipital alpha sources, indicating higher phase consistency. We also compared post-stimulus alpha power to rule out the possibility that wPPC differences were a result of less accurate phase estimation due to low alpha power (Cohen, 2014a). We observed a typical decrease in alpha power related to stimulus processing, but this effect did not differ between illusion and non-illusion trials.

**Fig 8.**
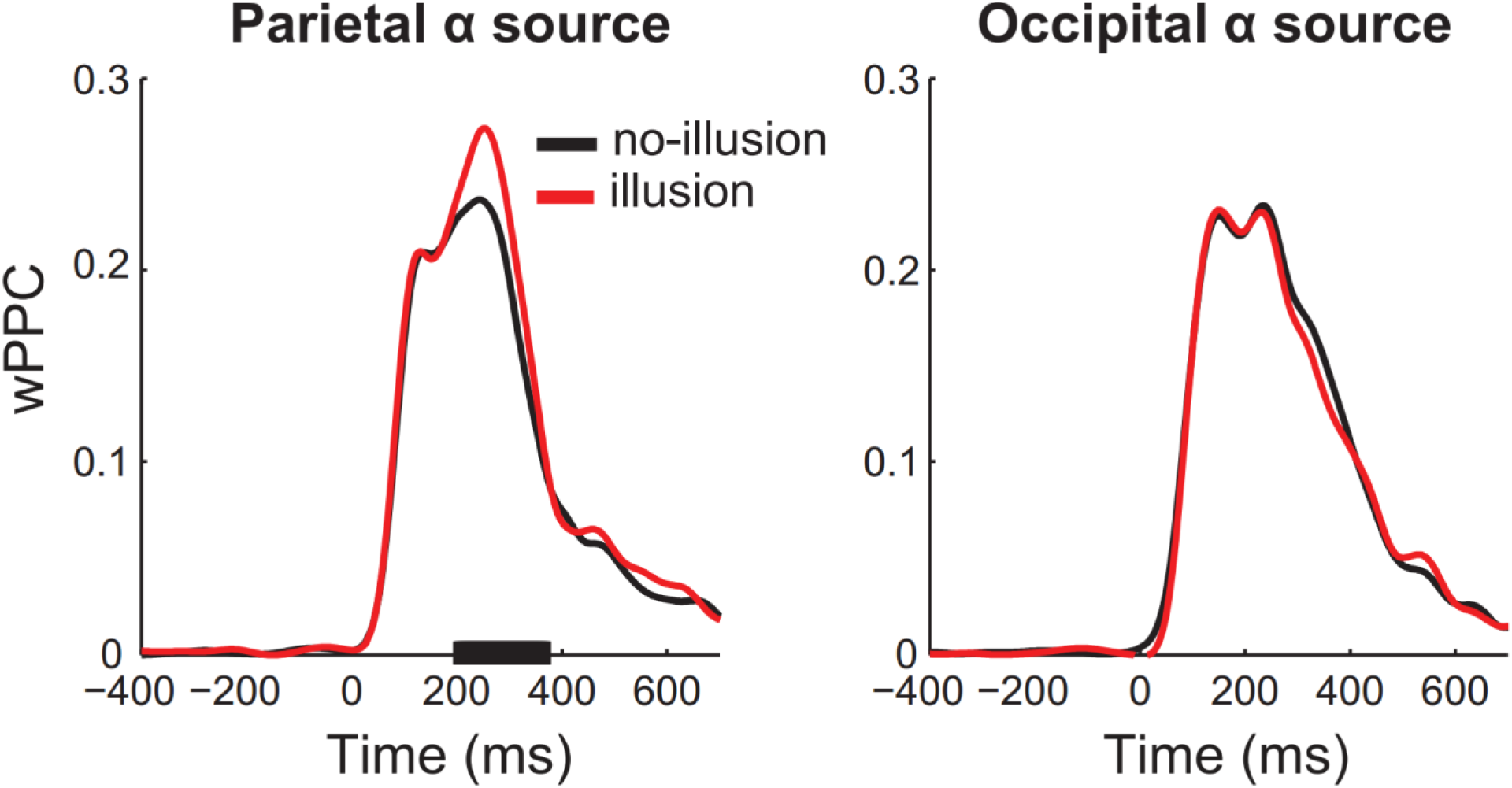
Pairwise phase consistency for illusion and non-illusion trials. Time-courses of weighted pairwise phase consistency (wPPC) at 11.43 Hz for illusion and non-illusion trials plotted separately for parietal (left panel) and occipital (right panel) alpha sources. Black bar on the x-axis represents time points at which wPPC for illusion vs. non-illusion trials was significantly different (corrected for multiple comparisons using cluster-based permutation testing).

## DISCUSSION

We validated and extended the original theoretical account of the triple-flash illusion proposed by Bowen (1989), according to which the illusory third-flash percept arises when the delay between the two veridical flashes matches the period of a hypothetical oscillatory impulse response function (IRF) generated in response to each stimulus. In Experiment 1, we demonstrated that the subject-specific inter-flash delay for which the illusory perception is maximized was strongly correlated with the period of the oscillatory IRF, which reverberates at ~10 Hz (Fig. 4C). The subject-specific IRF was derived from EEG responses to white-noise luminance sequences by cross-correlating the two signals (VanRullen and Macdonald, 2012). In Experiment 2, when fixing the inter-flash delay (87.5 ms) for all participants, we demonstrated that individual alpha peak frequency (IAF) of parietal but not occipital alpha sources was correlated with the overall proportion of illusory percepts: The closer participant’s parietal alpha peak was to 11.43 Hz (the 87.5 ms delay, expressed in Hz), the more illusions were perceived. Together, these results point to an active or ‘driving’ (as opposed to modulatory) role of alpha-band oscillations in perception, and reveal that alpha-band reverberations to a single stimulus have direct consequences on perception spanning several subsequent alpha cycles. Moreover, these findings emphasize the importance of using the “individual differences” approach when studying perceptual cycles and their associated oscillatory signatures.

Parietal and occipital alpha sources were estimated from ICA and single dipole fitting. Although ICs are often dipolar (Delorme et al., 2012), the anatomical dissociation of occipital vs. parietal sources should be interpreted cautiously, considering that dipole localization was based on 64-channel EEG using standard electrode locations, and a standard anatomical head model (although standard head models can provide reasonably high localization accuracy; Fuchs et al., 2002). However, a differential role of occipital vs. parietal alpha in perception has been reported previously and is consistent with our findings. In a discrimination task, for example, lower alpha power in parietal sources (BA 7) preceded correct trials, and was interpreted to reflect information gating from occipital to dorsal parietal areas controlled by top-down effects of attention (van Dijk et al., 2008). Importantly, this parietal alpha source was distinct from the occipital alpha source identified from the resting-state recordings. Relatedly, behavioral effects of 10 Hz rTMS in another visual discrimination task were observed only when stimulating parietal, but not occipital areas (Jaegle and Ro, 2014). Although this effect might be due to stronger entrainment effects in parietal compared to occipital rTMS, the proposed active role of parietal alpha in modulating visual representations in lower visual areas could account for the differential rTMS effects (Foxe and Snyder, 2011; Palva and Palva, 2011; Kwon et al., 2016). Pre-stimulus alpha phase in fronto-parietal areas has also been demonstrated to affect the connectivity between occipital and parietal areas, with certain pre-stimulus alpha phases associated with increased connectivity and better near-threshold stimulus detection (Hanslmayr et al., 2013).

Only two empirical studies of the triple-flash illusion, to our knowledge, have been conducted since Bowen’s original report; they compared the optimal SOA of the triple-flash illusion between healthy controls and schizophrenia patients (Norton et al., 2008; Chen et al., 2014). In both studies, the average SOA that maximized illusory perception was longer in the schizophrenia patient group (130-150 ms) than in the control group (90-110 ms). Following Bowen’s model, the authors speculated that such differences could result from temporal dilation of the IRF in schizophrenia patients. Considering both our between-subject correlation analyses results (Fig. 4–5), and previous reports of slower IAF in patients with chronic schizophrenia and schizophrenia symptoms (Canive et al., 1998; Harris et al., 2006), a slowing down of occipito-parietal alpha rhythms seems a conceivable explanation for the Norton et al. and Chen et al. findings.

Although Bowen’s theoretical model explains the mechanics behind the triple-flash illusion (Fig. 1B), it does not explain why at the subject-specific optimal SOA, an illusory third-flash is only perceived half of the time (45% on average). We hypothesized that this probabilistic nature of the illusion could be related to moment-to-moment fluctuations in occipito-parietal alpha phase and power that are known to be perceptually relevant (for a review, see Vanrullen et al., 2011). In the EEG experiment (Experiment 2), we found that trial-to-trial variability in perception of the illusion was indeed related to the pre-stimulus alpha phase at parietal but not occipital alpha sources, such that illusion and non-illusion trials were associated with opposite alpha phases (Fig. 6). Illusion trials were also preceded by significantly lower pre-stimulus alpha-band power at parietal alpha sources (Fig. 7). These findings are in accordance with previous reports linking relatively lower occipito-parietal alpha power and certain alpha phases to higher cortical excitability and better visual sensitivity (Romei et al., 2008; Mathewson et al., 2009; Dugue et al., 2011; Lange et al., 2013). Variations in occipito-parietal alpha power are related not only to veridical but also illusory perception: Illusory percepts are more frequent when occipito-parietal alpha power is low, both within-and between-subjects (VanRullen et al., 2006; Lange et al., 2014; Cecere et al., 2015).

Presentation of stimuli in phase with endogenous alpha oscillations results in stronger phase consistency across trials as compared to jittered stimulation (Thut et al., 2011; Spaak et al., 2014; Notbohm et al., 2016). In the post-stimulus period, we found higher phase consistency (as measured by wPPC) for illusion than non-illusion trials at 11.43 Hz (the two-flash trial SOA of 87.5 ms, translated into Hz) for parietal but not occipital alpha sources (Fig. 8). The wPPC differences between illusion and non-illusion trials, as well as pre-stimulus phase-opposition effects, appear complementary with Bowen’s theoretical model, which posits that illusory third-flash perception is associated with an enhancement of response amplitude resulting from phase-aligned oscillatory responses to each stimulus. Specifically, when the first stimulus appears in phase with ongoing pre-stimulus alpha oscillations (i.e., at the “good” pre-stimulus phase), there is no or relatively little phase re-alignment (Fellinger et al., 2011; Gruber et al., 2014); the second stimulus thus also arrives in phase with the ongoing oscillations and in phase with the oscillatory response generated to the first stimulus, resulting in high response amplitude and a third-flash percept. However, when the first stimulus arrives slightly or completely out of phase with the ongoing alpha oscillations (i.e., at the “bad” pre-stimulus phase), the resulting phase alignment in response to both stimuli is less precise (weaker wPPC), and thus only two flashes are perceived.

In conclusion, using the triple-flash illusion – a third illusory flash perception when only two veridical ones are presented, separated by ~100 ms – we demonstrate that alpha-band oscillations not only modulate perception but have a driving impact, which can make one perceive something that is not there.

## ACKNOWLEDGEMENTS

This research was funded by the ERC grant P-CYCLES (N°614244) awarded to Rufin VanRullen.

